# Fungal-bacterial interactions enhance bacterial but constrain fungal resistance and recovery to extreme heat across soil moisture levels

**DOI:** 10.64898/2026.11.09.765442

**Authors:** Jingjing Shi, Armelle Vallat, Pilar Junier, Madhav P. Thakur

## Abstract

Soil fungi and bacteria are key drivers of nutrient cycling and plant productivity in terrestrial ecosystems, yet their responses to climate extremes are rarely studied as interacting partners rather than in isolation. Moreover, it remains largely unknown how fungal-bacterial interactions regulate fungal and bacterial stability under climate extremes, particularly resistance (immediate responses to climate extremes) and recovery (responses after climate extremes cease). Here, we conducted a soil microcosm experiment with a bacterial species (*Pseudomonas putida* KT2440) paired with one of four *Trichoderma* species (*T. hamatum*, *T. atroviride*, *T. koningiopsis*, and *T. harzianum*). These fungal-bacterial pairs were then exposed to a 5-day heat event (40°C/32°C, day/night) followed by a 14-day recovery period (25°C/17°C, day/night), at two soil moisture levels (30% and 5% gravimetric water content). Our results show that fungal-bacterial interactions asymmetrically affect fungal and bacterial responses to extreme heat. *Trichoderma* species facilitated the resistance and recovery of *P. putida* density (CFUs) to extreme heat, whereas *P. putida* constrained the resistance and recovery of fungal biomass (ergosterol) to extreme heat, regardless of soil moisture conditions. Moreover, faster-growing *Trichoderma* species were associated with greater facilitation of bacterial resistance and recovery. Bacterial density also covaried positively with fungal biomass during heat stress under both soil moisture conditions, suggesting that net fungal facilitation enhances bacterial persistence under extreme heat. Together, our findings reveal that fungal-bacterial interactions can stabilize one microbial partner while increasing the vulnerability of the other, highlighting the contrasting consequences of biotic interactions under climate extremes.

## 1. Introduction

Soil microorganisms, particularly fungi and bacteria, drive key ecosystem processes such as nutrient cycling, carbon sequestration, and plant productivity in terrestrial environments (Wei et al., 2019; Sokol et al., 2022; Tao et al., 2023). Climate extremes, such as extreme heat and drought, are increasing in frequency and intensity (IPCC, 2023; Sanches et al., 2023), threatening both the stability and functionality of soil microbial communities (Bardgett and Caruso, 2020; Jansson and Hofmockel, 2020; Philipp et al., 2025). The stability of soil fungi and bacteria under climate extremes can be characterized by their resistance (immediate responses to climate extremes) and recovery (responses following the cessation of climate extremes) (Bardgett and Caruso, 2020; Thakur et al., 2022). Although the responses of soil fungi and bacteria to climate extremes have been widely studied (Ochoa-Hueso et al., 2018; Bei et al., 2023; Cordero et al., 2023; Knight et al., 2024; Sun et al., 2025), how fungal-bacterial interactions shape microbial resistance and recovery under climate extremes remains a major research gap (Shi and Thakur, 2023).

Fungal-bacterial interactions play important roles in biogeochemical cycling and plant health and can be mediated by diverse physical, chemical, and molecular mechanisms (Deveau et al., 2018; Zhou et al., 2022; Yusuf et al., 2025). For example, fungal hyphae can act as highways for bacterial dispersal, and hyphal exudates can serve as sources of carbon and nutrients, while bacteria can promote or suppress fungal growth through specific metabolites (Kohlmeier et al., 2005; Deveau et al., 2010; Warmink et al., 2011; Richter et al., 2024; Vieira et al., 2025). Depending on the partners involved, these mechanisms produce outcomes ranging from antagonism to mutualism (Deveau et al., 2018; Zhou et al., 2022; Wang and Kuzyakov, 2024). Abiotic environmental stress can strengthen or weaken existing positive or negative fungal-bacterial interactions, or shift them between negative and positive outcomes (Velez et al., 2018; Gao et al., 2022). In turn, these interactions may influence how each partner responds to stress, as fungi and bacteria can modify each other’s microenvironment and resource supply (Nazir et al., 2014; Worrich et al., 2017; Deveau et al., 2018).

Fungi and other hyphal organisms can promote bacterial survival and growth under abiotic stress (Worrich et al., 2017; Yang et al., 2018; Venkatesh et al., 2022). For instance, the mycelium of the oomycete *Pythium ultimum* facilitated spore germination and vegetative growth of *Bacillus subtilis* in dry, oligotrophic environments by transferring water and nutrients (Worrich et al., 2017). The soil saprotrophic fungus *Lyophyllum* sp. strain Karsten increased soil pH, thereby promoting the survival of *Paraburkholderia terrae* BS001 under acidic conditions (Yang et al., 2018). Furthermore, *Aspergillus flavus* enhanced *Ralstonia solanacearum* survival under nutrient starvation and cold stress by allowing the bacterium to colonize its chlamydospores (Venkatesh et al., 2022). Such mechanisms may help bacteria withstand stressful conditions. In contrast, evidence for bacterial effects on fungal stress tolerance is limited and inconsistent. *Burkholderia terrae* BS001 (reclassified as *Paraburkholderia terrae* BS001) protected *Lyophyllum sp.* strain Karsten against the antifungal agent cycloheximide (Nazir et al., 2014). Besides, endobacteria modulate fungal resistance to heavy metals in species-specific ways, either enhancing or reducing tolerance depending on the pairing of the fungus, the bacterial symbiont, and the specific metal (Lupini et al., 2022). Together, these studies suggest that fungal-bacterial interactions can shape microbial fitness under abiotic stress, but how they influence microbial resistance and recovery under extreme heat, particularly for fungi, remains unclear.

To investigate how fungal-bacterial interactions influence microbial stability (resistance and recovery) under extreme heat, we conducted a soil microcosm experiment using pairwise combinations of several *Trichoderma* spp. and *Pseudomonas putida* as a simplified fungal-bacterial interaction model system. These genera are widespread in soils and are often applied together in biocontrol and plant growth promotion (Harman et al., 2004; Volke et al., 2020; Poveda and Eugui, 2022; Cejas et al., 2026). We tested how fungal biomass (quantified by ergosterol analysis) and bacterial density (quantified by colony-forming units assay) respond to an extreme heat event under contrasting soil moisture conditions, with and without the interacting partner. We hypothesize that (1) *Trichoderma* will enhance the resistance and recovery of *P. putida* to extreme heat, given previous evidence that fungi can protect bacteria under abiotic stress; (2) *P. putida* will alter the resistance and recovery of *Trichoderma* to extreme heat, but the direction of this effect is uncertain. Bacteria could enhance fungal resistance and recovery by protecting fungi from stress or reduce them through resource competition or inhibitory metabolites. Finally, we explored whether these interaction effects differed between high (30%) and low (5%) soil moisture.

## 2. Materials and Methods

### 2.1 Bacteria and fungi

The bacterial strain used in our study was *Pseudomonas putida* KT2440 (GFP-tagged) (Dechesne et al., 2010; Pion et al., 2013; de Lorenzo et al., 2024), obtained from the Laboratory of Microbiology (Institute of Biology, University of Neuchâtel). The two fungal strains, *Trichoderma atroviride* NEUM134 and *Trichoderma harzianum* NEUM136, were also obtained from the Laboratory of Microbiology (Institute of Biology, University of Neuchâtel). The other two fungal strains were obtained from Westerdijk Fungal Biodiversity Institute (Utrecht, Netherlands): *Trichoderma hamatum* CBS 121.13 and *Trichoderma koningiopsis* CBS 132570. *Pseudomonas putida* KT2440 was grown at 28 °C on LB medium (LB-Lennox, Carl Roth, Germany; 10g/L tryptone, 5 g/L yeast extract, and 5 g/L sodium chloride) supplemented with 1.5% agar if required. *Trichoderma* species were grown at 20 °C on PDA (Carl Roth, Germany; 4 g/L potato infusion, 20 g/L glucose, and 15 g/L agar) for general growth and maintenance.

### 2.2 Soil microcosm experiment

A universal gardening soil (Profi Universalerde, Terre Suisse AG, Switzerland) was sieved through a 2 mm stainless-steel mesh, then mixed with sand (soil: sand ratio = 3:1, w:w). The mixed soil was oven-dried at 70 °C for 72 hours, then autoclaved three times (121 °C for 25 min) to ensure optimal sterilization. The moisture of the sterilized soil was then adjusted to the target gravimetric water content (GWC) of 30% and 5% by adding a calculated amount of autoclaved deionized water. Finally, 30 g of prepared soil was added to each microcosm (150 mL screw-cap polypropylene jar).

The fungi were cultivated on PDA at 20°C for inoculation. Fungal plugs were prepared from the actively growing edge of the fungal culture plates with a stainless-steel cork borer (diameter: 5 mm), and 2 fungal plugs were added into the soil together at the centre of each soil microcosm. To prepare the bacterial inoculum, *P. putida* was cultivated in LB broth at 28 °C and 220 rpm for 6 hours. We then centrifuged the liquid culture (3900 rpm for 6 minutes) to pellet the bacterial cells, resuspended them, and washed them twice with sterile deionized water. We resuspended the washed bacterial cells in sterile deionized water, measured the optical density at 600 nm (OD600) with a photometer (DEN-600, Biosan, Latvia), and adjusted it to a final OD600 of 0.2 with sterile deionized water for inoculation. We added 800 µL of bacterial inoculum to the soil surface of each microcosm, pipetting in a clockwise spiral from the center to the edge. We then placed the soil microcosms in incubators (HPP410eco, Memmert, Germany) for climatic treatments (details below). We processed the sterilized soil, prepared fungal and bacterial inocula, and inoculated fungi and bacteria under aseptic conditions in a biosafety cabinet (FlowGard^TM^, BioTool, Switzerland).

Experimental treatments were applied as follows: two levels of temperature (ambient temperature: 25°C for 15 hours and 17°C for 7 hours for the day/night cycle; extreme heat: 40°C for 15 hours and 32°C for 7 hours for the day/night cycle; 1 hour ramping time between temperature shifts); two levels of soil moisture (30% and 5% GWC); two levels of bacteria inoculation (non-bacteria; *P. putida*); 5 levels of fungal inoculation (non-fungi; *T. hamatum*; *T. atroviride*; *T. koningiopsis*; *T. harzianum*). We destructively harvested microcosms to estimate bacterial density and fungal biomass (details below) on day 1 (the day after the experiment started), day 7, day 12, and day 26. The heat treatment was applied from day 7 to day 12, after which temperatures returned to ambient conditions until day 26; accordingly, day 12 represents the end of the heat-stress period (resistance phase) and day 26 represents the end of the post-stress recovery period (recovery phase). Each treatment combination was replicated five times (except three times for the first and second harvests on day 1 and day 7, respectively), resulting in a total of 520 microcosms.

### 2.3 Measurements

To measure bacterial density in soil, we took a 1 g fresh soil subsample from the microcosm after gently homogenizing it with a stainless-steel spatula (flame-sterilized and cooled before use), transferred it into a 15 mL centrifuge tube (Greiner Bio-One, Austria), and added 9 mL of sterile 1× PBS. The samples were stored at 4°C (standing in racks) for the CFUs assay, with storage times up to 4 weeks (Lee et al., 2021). The CFUs assay was conducted using the spread plate method (Sanders, 2012). Specifically, soil samples were vortexed vigorously at full intensity for 5 minutes with a vortex mixer (Vortex-Genie 2, Scientific Industries, USA) to detach bacterial cells. After that, a 10-fold dilution series of the soil slurry was prepared directly in a 96 Deep-well plate with sterile 1 × PBS, and 100 µL aliquots of each dilution were added to an LB plate and spread with 10-15 sterile glass beads (diameter: 3 mm) by horizontal shaking to allow the beads to run in random directions. After removing the glass beads, the plates were sealed with Parafilm, incubated at 28°C for 12-15 hours, and then counted for bacterial colonies.

To measure fungal biomass in soil, we took a 1 g fresh soil subsample from the homogenized soil in the microcosm, transferred it to a 15 mL centrifuge tube (Greiner Bio-One, Austria), and added 4 mL of 10% KOH in methanol (Supelco, Germany). The samples were stored at-20°C (standing in racks) before ergosterol extraction. We used the alkaline extraction method (Bååth, 2001) to extract ergosterol from soil samples. More specifically, we first sonicated samples in tubes for 15 min, then placed them in a water bath at 70 °C for 90 min. Then, we added 1 mL of Milli-Q water and 2 mL of n-Hexane (Supelco, Germany) to the tube. We then closed the tube, vortexed it for 30 s on a vortex mixer, and centrifuged it at 3900 rpm for 12 min. After centrifugation, we transferred 1 mL of the top phase into a 5 mL glass test tube (Duran, Germany). We then repeated this by adding 1 mL of n-Hexane (Supelco, Germany) to the 15 mL tubes, vortexing, and centrifuging to obtain another 1 mL of the top phase, which we transferred again to the glass test tube. To evaporate the hexane, we placed the glass test tubes in a water bath at 50 °C in a chemical fume hood overnight. Afterward, we added 1 mL methanol (Supelco, Germany) to each test tube and sealed the tubes with a glass bead (diameter: 10 mm). We then mixed the ergosterol extracts with methanol for 30 secs on a vortex mixer, sonicated the tubes for 4 min, and mixed again for 30 secs on a vortex mixer. Finally, we filtered the solution from the test tubes into a 2 mL amber vial (A-Line certified screw-top vial, Agilent, USA) using a 0.45 µm PTFE syringe filter (Avantor, USA). The ergosterol concentration in the extracts was then determined by QTRAP 6500+ LC-MS/MS system (Sciex, Marlborough, MA). A 1 μl aliquot of the sample was injected onto an ACQUITY UPLC® BEH C18 column (2.1 x 50 mm, 1.7 µm) (Waters, Milford, MA). The flow rate and column temperature were kept at 0.400 ml/min and 30 °C, respectively. Mobile phase A was Milli-Q water, and mobile phase B was methanol (ULC-MS grade, Biosolve, Valkenswaard, The Netherlands). The following gradient was used: 80-100% B in 5 min, holding at 100% for 1.7 min, followed by re-equilibration at 80% B for 1.3 min, for a run time of 8 min. Mass spectrometry detection was performed in multiple reaction monitoring (MRM) mode in APCI positive ionization, with the transition 379.5/69.4. Quantification was done by external calibration using a linear regression from seven calibration points (25 to 2500 ng/mL).

### 2.5 Statistics

All statistical analyses were conducted in R version 4.5.0 (R Core Team, 2025). To test the effects of temperature treatment, soil moisture, fungal treatment (either as two levels: fungal presence/absence or as five levels to include all fungal species), and time on bacterial density, we used linear regressions (with normal distribution). Models initially included all main effects and their interactions. Model complexity was reduced by sequentially removing higher-order interactions and comparing nested models using likelihood ratio tests (ANOVA). Time points were additionally incorporated into the model as a quadratic term (poly(time_day, 2)) to allow non-linear temporal dynamics in bacterial density, as shown in Figure S2. Three candidate models: full interactions model with time as a linear term, reduced interactions model with time as a linear term, and reduced interactions model with time as a quadratic term, were compared using the model.sel () function from the ‘MuMIn’ package (Bartoń, 2023). We compared candidate models using the small-sample-corrected Akaike information criterion (AICc), which accounts for model complexity and provides a correction for finite sample sizes (Burnham and Anderson, 2002). We analyzed fungal biomass using the same approach, with temperature treatment, soil moisture, bacterial presence, fungal species, and time as main effects. The model with the lowest AICc was selected for subsequent analyses, and the candidate model comparisons for bacterial density and fungal biomass are provided in the supplementary information (Tables S1-S4).

Prior to analysis, bacterial density (CFUs) was log₁₀-transformed, and fungal biomass (ergosterol) was ln-transformed to satisfy assumptions of normality and homogeneity of variance. Following model selection, residual plots were inspected visually to confirm that these assumptions were met. The significance of main effects and interactions was assessed using Type III analyses of variance, implemented with the Anova() function from the ‘car’ package (Fox and Weisberg, 2019). The statistical results are presented in supplementary information (Table S5-S6).

To quantify the magnitude of heat-treatment effects on bacterial density and fungal biomass, we calculated standardized effect sizes (Cohen’s d) for sampling points at the resistance phase (day 12) and recovery phase (day 26). Estimated marginal means (EMMs) were derived from previously selected linear models using the’emmeans’ package (Lenth, 2017). Pairwise contrasts between heat and ambient temperature treatments were extracted separately for each combination of soil moisture level and inoculation treatment at the resistance sampling point and at the recovery sampling point. Because models were fitted to log-transformed data, contrasts represent differences on the log scale; Cohen’s d and its 95% confidence intervals were therefore computed as standardized effects on the log scale using the eff_size() function, which divides each contrast by the model’s pooled residual standard deviation.

Finally, we explored the relationship between bacterial density and fungal biomass during the resistance and recovery phases using Pearson’s correlation. Before analysis, we evaluated the assumptions of bivariate normality, linearity, and homoscedasticity using diagnostic plots and confirmed they were satisfactorily met.

## 3. Results

### 3.1 Bacterial and fungal responses to extreme heat

Fungal presence consistently increased both the resistance and recovery of bacterial density to extreme heat under both soil moisture conditions (Figure 1A). The negative effects of extreme heat on bacterial density during and after the heat period were more pronounced under high soil moisture (30% GWC) than under drought (5% GWC), regardless of fungal presence (Figure 1A). In contrast, bacteria reduced both fungal resistance and recovery from extreme heat under both moisture conditions (Figure 1B). Under high soil moisture (30% GWC), extreme heat significantly reduced fungal biomass regardless of bacterial presence, whereas under drought (5% GWC), bacteria were required for this reduction. After the heat period, fungal biomass showed under-recovery only in drought soil when bacteria were present (Figure 1B).

**Figure 1.**
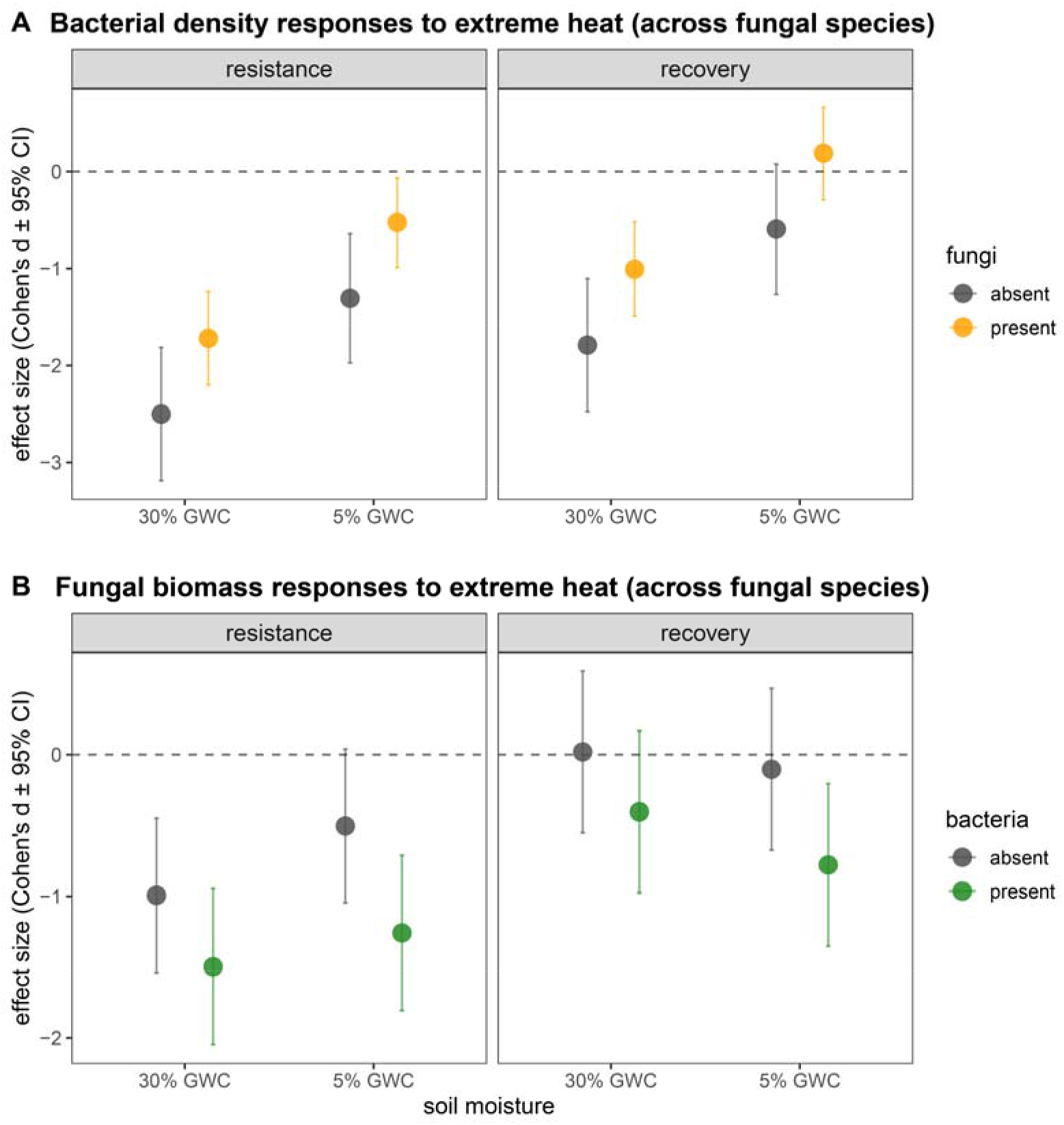
Bacterial and fungal responses to extreme heat (across fungal species). The resistance and recovery of bacteria (A) and fungi (B) to extreme heat under two soil moisture conditions. The effect sizes are statistically significant when their 95% confidence intervals (CI) do not overlap with zero. GWC: Gravimetric water content.

Furthermore, bacterial resistance and recovery to extreme heat varied among fungal species and generally followed the rank order of fungal growth rates (Figure S1), with *T. harzianum* (fastest among the four fungi) showing the most facilitative effects and *T. hamatum* (slowest among the four fungi) the least facilitative effects under both soil moisture conditions (Figure 2). Conversely, all fungal species generally exhibited reduced resistance and recovery in the presence of bacteria under both soil moisture conditions, with *T. hamatum* showing the highest decline (Figure 3).

**Figure 2.**
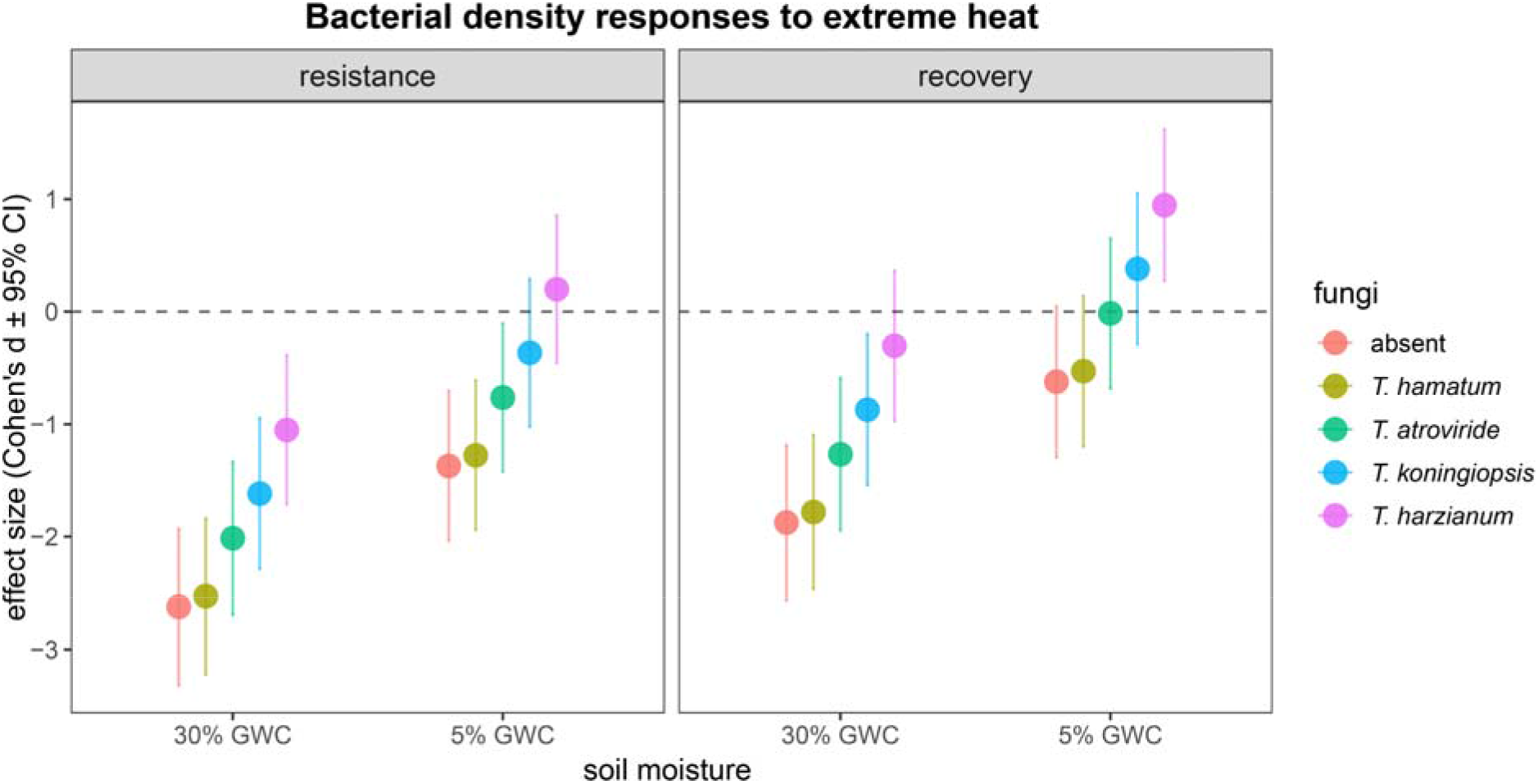
Bacterial density responses to extreme heat in the absence of fungi and in the presence of different fungal species. The effect sizes are statistically significant when their 95% confidence intervals (CI) do not overlap with zero. GWC: Gravimetric water content.

**Figure 3.**
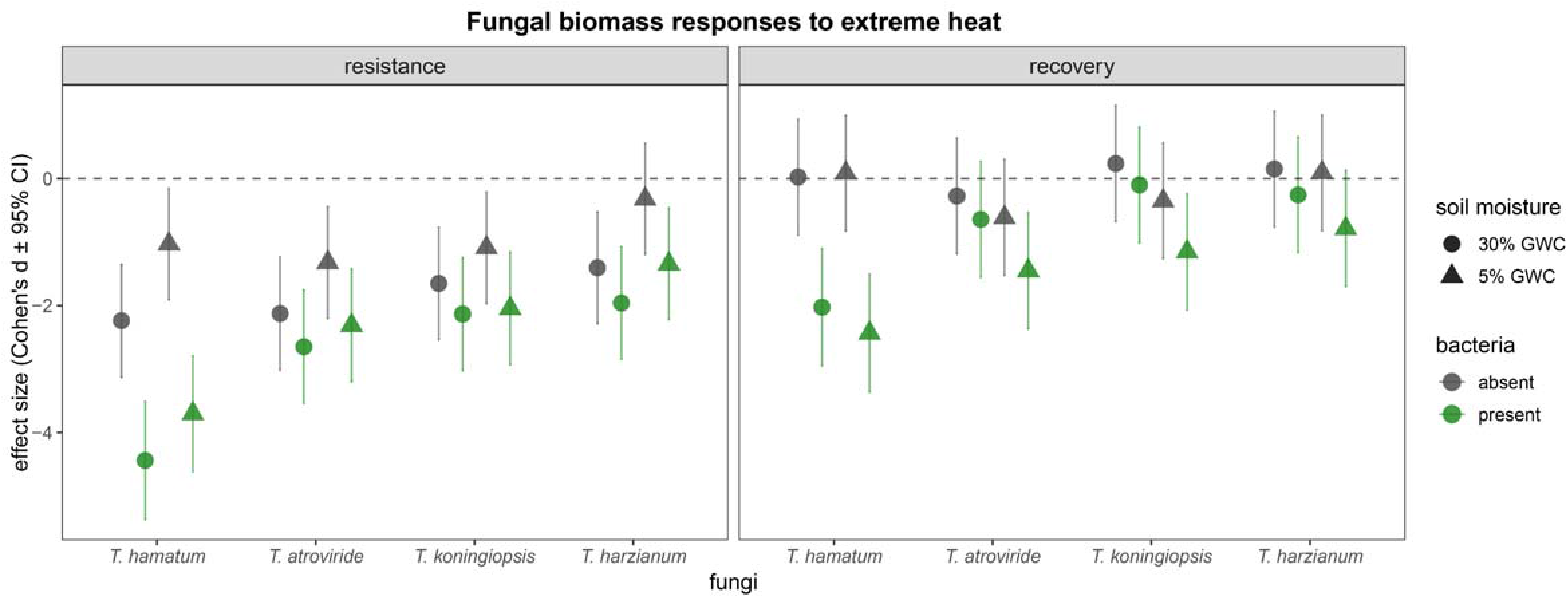
Fungal biomass responses to extreme heat (fungal species level) in the absence and presence of bacteria. The effect sizes are statistically significant when their 95% confidence intervals (CI) do not overlap with zero. GWC: Gravimetric water content.

### 3.2 Bacterial density and fungal biomass dynamics in soil

Bacterial density varied across the experimental period with soil moisture, temperature treatment, and fungal inoculation (Table S5). Bacterial density was higher under high soil moisture (30% GWC) than under drought (5% GWC) and was lower under extreme heat than under ambient temperature (Figure S2). Bacterial density also differed among fungal inoculation treatments. Temperature effects on bacterial density depended on soil moisture and fungal inoculation, as shown by significant temperature × soil moisture and temperature × fungi interactions (Table S5). Temporal trajectories of bacterial density further differed among soil moisture, temperature, and fungal inoculation treatments, as indicated by significant interactions between time and each of these factors (Table S5).

Fungal biomass also varied across the full experimental period with soil moisture, temperature treatment, bacterial inoculation, and fungal species identity (Table S6). Fungal biomass was higher under high soil moisture than under drought, lower under extreme heat than under ambient temperature, and lower in the presence of *P. putida* than in the absence of bacteria (Figure S3). Fungal biomass also differed among fungal species. Temperature effects on fungal biomass depended on bacterial inoculation, soil moisture, and fungal species identity, as shown by significant temperature × bacteria, temperature × soil moisture, and temperature × fungal species interactions (Table S6). The effect of bacterial inoculation on fungal biomass also depended on soil moisture and fungal species identity, and fungal biomass trajectories varied among treatment combinations over time, as indicated by higher-order interactions among these variables (Table S6).

### 3.3 Correlation between bacterial density and fungal biomass in soil

At the end of the resistance phase (day 12), correlations between bacterial density and fungal biomass were significantly positive in heat-exposed treatments but nonsignificant in ambient-temperature treatments, under both soil moisture conditions (Figure 4A). At the end of the recovery phase (day 26), bacterial density in ambient temperature treatments was negatively correlated with fungal biomass under high soil moisture (30% GWC), whereas no significant correlations between bacterial density and fungal biomass were found in other combinations of soil moisture and temperature treatments (Figure 4B).

**Figure 4.**
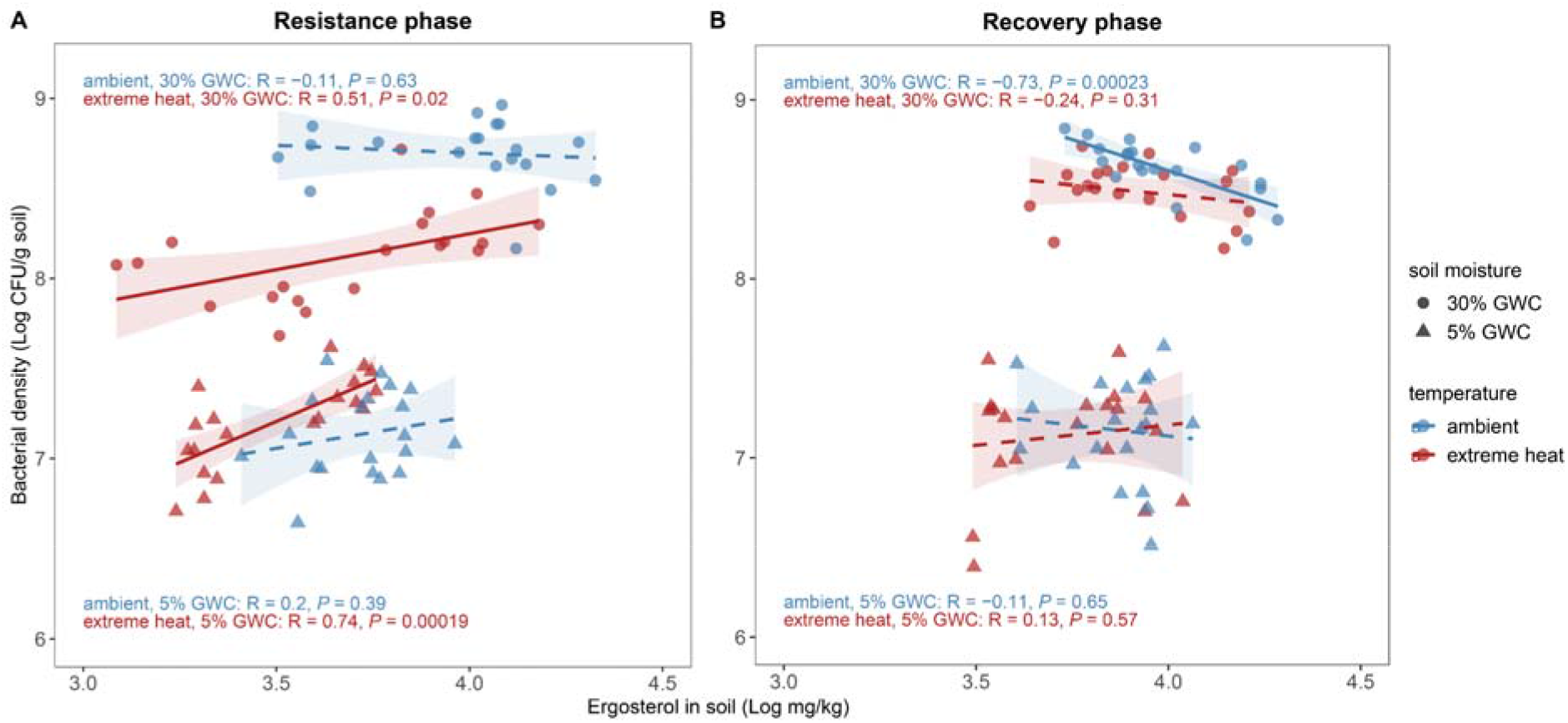
Correlation between fungal biomass and bacterial density. Pearson’s correlation between fungal biomass (log_10_-transformed) and bacterial density (log_10_-transformed) at day 12 (resistance sampling point) (A) and day 26 (recovery sampling point) (B). Solid lines indicate significant correlations, and dashed lines represent non-significant correlations. GWC: Gravimetric water content.

## 4. Discussion

Microbial interactions fundamentally drive microbial stability and ecosystem functioning, yet their role in microbial responses to climate extremes remains poorly understood (Wagg et al., 2019; Ratzke et al., 2020; Thakur et al., 2022; Shi and Thakur, 2023). Using a soil microcosm experiment and a simplified fungal-bacterial interaction model system, we explored how cross-kingdom interactions influence microbial stability under extreme climatic events such as extreme heat. We found that the presence of fungi (*Trichoderma*) facilitated the resistance and recovery of bacteria (*P. putida*) to extreme heat, whereas the presence of bacteria generally impeded the resistance and recovery of fungi to extreme heat, regardless of soil moisture conditions. These findings highlight that fungal-bacterial interactions can shape each other’s stability under climate extremes asymmetrically, with outcomes differing between the fungal and bacterial partners.

In our study, *Trichoderma* species mitigated heat-induced declines in *P. putida* density during resistance and recovery phases, under both high soil moisture (30% GWC) and drought (5% GWC). Our results align with previous findings about the protective effects of fungi on bacteria under abiotic stress (Worrich et al., 2017; Hestrin et al., 2022; Pathan et al., 2023). Moreover, *Trichoderma* species with higher growth rates were associated with stronger facilitative effects on bacterial resistance and recovery, and bacterial density covaried positively with fungal biomass during the heat event, suggesting that fungal facilitation depends on the extent of hyphal growth and may involve hyphal-mediated processes. For instance, fungal hyphae may enhance bacterial stress tolerance by enabling bacteria to colonize or migrate along hyphae to microhabitats with more favorable water and nutrient conditions (Nazir et al., 2010; Vieira et al., 2025). We speculate that fast-growing fungi may develop more extensive hyphal networks, which could enhance bacterial dispersal along fungal highways and potentially provide microhabitats that buffer bacteria against heat stress. In addition, bacteria can produce extracellular polymeric substances (EPS) that form biofilms on the surfaces of fungal hyphae, thereby protecting the cells from environmental stress (Miquel Guennoc et al., 2018; Kjeldgaard et al., 2019; Xie et al., 2024). Although bacterial dispersal and biofilm formation were not specifically measured in our soil microcosm experiment, we observed *P. putida* colonies dispersing onto the hyphae of *Trichoderma* and biofilms forming within the hyphal network of *Trichoderma* on agar plates, as revealed by fluorescence and scanning electron microscopic images (Figure 5). Future studies could test whether hypha-associated bacterial dispersal and biofilm formation also occur in soil and whether these processes contribute to enhanced bacterial resistance and recovery under extreme heat and drought.

**Figure 5.**
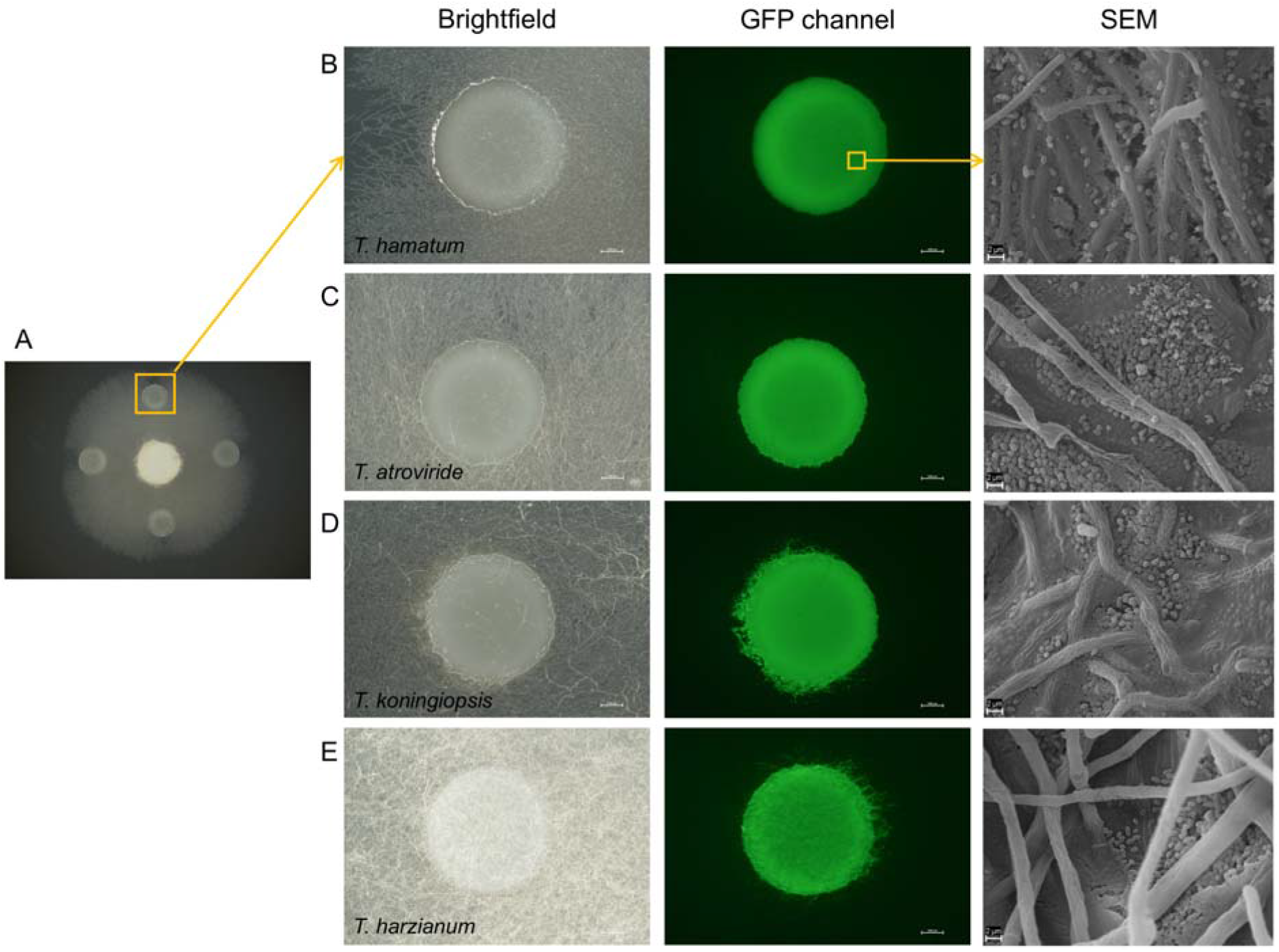
Microscopic images of *P. putida* and *Trichoderma*. (A) Bacterial colonies around the fungal growth center (agar plug) in the PAD plate under the stereomicroscope Nikon SMZ 25. *P. putida* colony dispersal on the hyphal network of *T. hamatum* (B), *T. atroviride* (C), *T. koningiopsis* (D), and *T. harzianum* (E) in brightfield and the GFP channel of fluorescence microscopy with Nikon SMZ 25 (scale bar: 1000 µm), and scanning electron microscopy with Zeiss EVO 40 (scale bar: 2 µm), from left to right, respectively. Supplementary information provides details of sample preparation for fluorescence microscopy and scanning electron microscopy.

Conversely, *P. putida* exhibited negative effects on the resistance and recovery of *Trichoderma* to extreme heat under both soil moisture levels. Among the four *Trichoderma* species, *T. hamatum* showed the greatest decline in both resistance and recovery under extreme heat when *P. putida* was present. Previous studies have reported stimulatory effects of *P. putida* on the growth of *T. aggressivum* and inhibitory effects of rhizosphere-isolated *Pseudomonas* on the growth of *T. harzianum* and *T. virens* (Li et al., 2019; Kosanovic et al., 2020), suggesting that the effects of *Pseudomonas* on *Trichoderma* are likely to be both positive and negative depending on species identity. *P. putida* was associated with lower *Trichoderma* biomass and more negative fungal responses to extreme heat, indicating that bacterial presence may have constrained fungal resistance and recovery. Whether these effects arose from direct bacterial inhibition, competition for resources, altered fungal physiology, or heat-induced changes in bacterial-fungal interactions remains unresolved and warrants further investigation.

The positive correlations between bacterial density and fungal biomass occurred during extreme heat (resistance phase) and were stronger under low rather than high soil moisture. Harsher environmental conditions may favor positive interactions between fungi and bacteria according to the stress gradient hypothesis (Hammarlund and Harcombe, 2019). For instance, positive correlations in co-occurrence networks of fungi and bacteria increased under drought in a soil experiment (Gao et al., 2022). Besides, fungal-bacterial interactions shifted from competition to cooperation as nutrient limitation increased (Velez et al., 2018). After the extreme heat ended (recovery phase), correlations between bacterial density and fungal biomass became nonsignificant in heat-exposed treatments, whereas non-heat-exposed treatments showed significantly negative correlations between bacterial density and fungal biomass under high soil moisture, suggesting potentially increased competition between bacteria and fungi for soil resources (Wang and Kuzyakov, 2024). Together, these findings suggest that climate extremes can modulate fungal-bacterial interaction dynamics.

Our study has several limitations. First, we used a single bacterial species and four species of one fungal genus in a simplified pairwise system. The asymmetric effects we observed may therefore not generalize to other fungal-bacterial combinations or to diverse soil communities, where additional competitors, predators, and mutualists could modify both directions of the interaction. Second, we focused on population-level responses and did not measure the mechanisms underlying the observed interactions in the soil. Our microscopy observations of bacterial dispersal along hyphae and biofilm formation came from agar plates, so whether such physical interactions explain fungal facilitation of bacteria in soils remains to be tested, for example with microfluidic soil-chip systems (Mafla-Endara et al., 2021). Likewise, measuring stress-tolerance traits such as dormancy and sporulation, and molecular responses such as heat shock protein expression, would help understand the underlying mechanisms of microbial responses to extreme events (Bérard et al., 2015; Shade, 2023; Knight et al., 2024). Finally, we imposed a single heat event at two constant moisture levels and followed recovery for 14 days. Natural climate extremes vary in intensity, duration, and frequency, and may recur or co-occur. Moreover, soil structure, the spatial distribution of water and nutrients, pH, and plant roots can all influence microbial interactions under climate extremes (Custódio et al., 2022). Incorporating these factors in future experiments would clarify how fungal-bacterial interactions shape microbial stability under climate extremes in natural soils.

## 5. Conclusions

Our results reveal asymmetric effects of fungal-bacterial interactions on microbial responses to extreme heat. *Trichoderma* enhanced both the resistance and recovery of *P. putida* density, whereas *P. putida* constrained both the resistance and recovery of *Trichoderma* biomass, and these effects were consistent across contrasting soil moisture conditions. Faster-growing *Trichoderma* species provided greater benefits to bacteria, suggesting that fungal growth traits shape the strength of facilitation. This asymmetry indicates that the direction and magnitude of fungal-bacterial interactions under climate extremes cannot be assumed to be reciprocal, which should be considered when predicting microbial responses to climate change. Identifying the functional traits and mechanisms that underlie this asymmetry, and testing whether it holds in more diverse communities and under recurrent or compound extremes, are key next steps toward a mechanistic understanding of soil microbial stability under climate change.

## Acknowledgements

We are thankful for the support from the Swiss National Science Foundation (awarded to MPT, grant no. 310030_212550). We thank Roem Yesil, Mailén Guerra Moreno, and Anine Wyser for help with experimental setup and sampling. We thank Desirée Dürr and Marek Kaminek for assistance with the scanning electron microscopy.

## Author Contributions

**JS**: conceptualization, investigation, methodology, data curation, formal analysis, writing-original draft, writing-review & editing; **AV**: methodology, data curation, writing-review & editing; **PJ**: conceptualization, supervision, resources, writing-review & editing; **MPT**: conceptualization, formal analysis, funding acquisition, supervision, resources, writing-original draft, writing-review & editing.

## Data Availability Statement

Data are available from the Figshare Repository: https://doi.org/10.6084/m9.figshare.34962364

