## Supplementary information for "Fungal-bacterial interactions enhance bacterial but constrain fungal resistance and recovery to extreme heat across soil moisture levels"

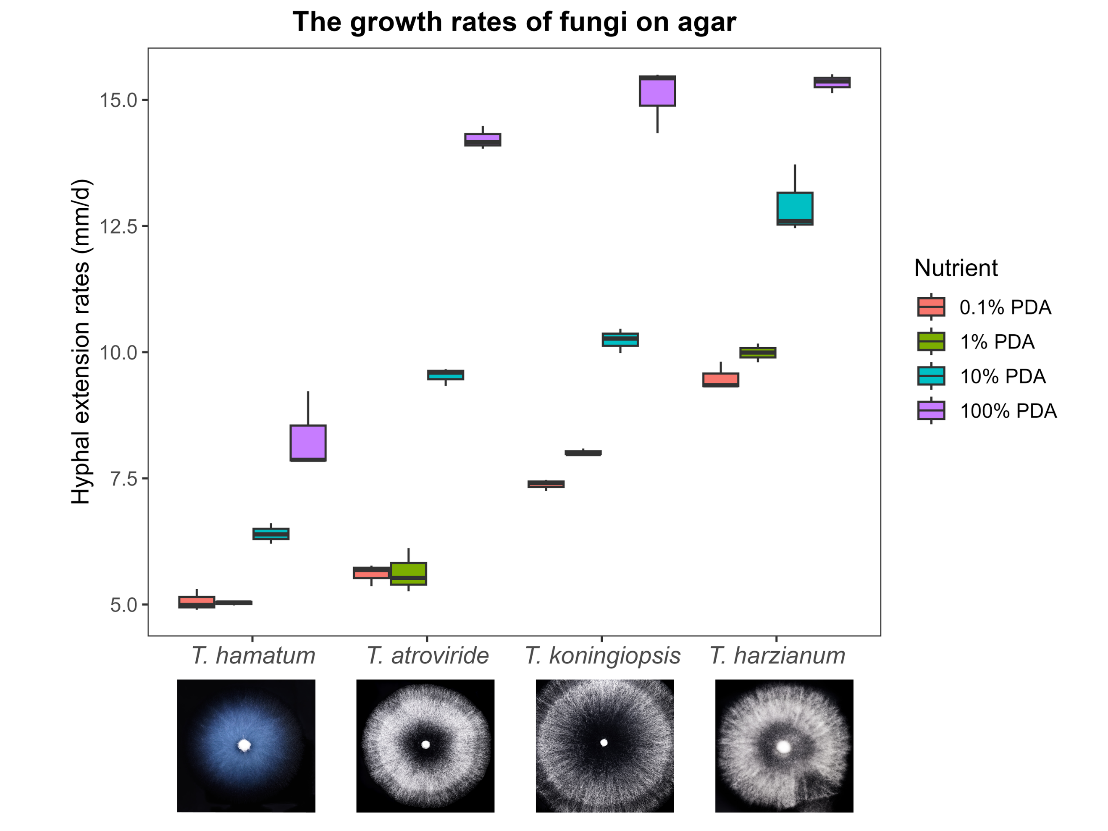

**Figure S1. Fungal growth rates on agar.** Hyphal extension rates of the *Trichoderma* species on PDA with 4 nutrient levels. Images of each fungus on PDA plates that were supplemented with charcoal (4g/L PDA) for a black background contrast (photo credits: Nicolas Stettler).

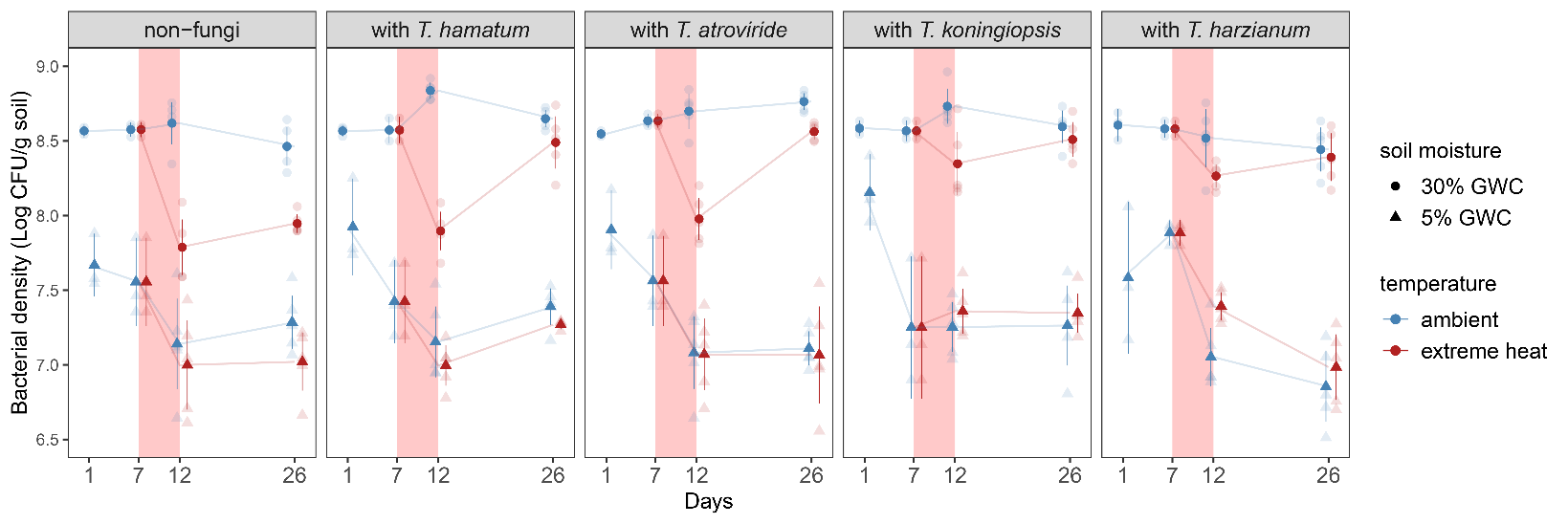

**Figure S2. Bacterial density dynamics in soil.** Bacterial density (log_10_-transformed) without fungi and with the presence of different *Trichoderma* species (mean ± standard error, n = 3 for day 1 and 7; n = 5 for day 12 and 26). The red region between day 7 and day 12 indicates the period of extreme heat. GWC: Gravimetric water content.

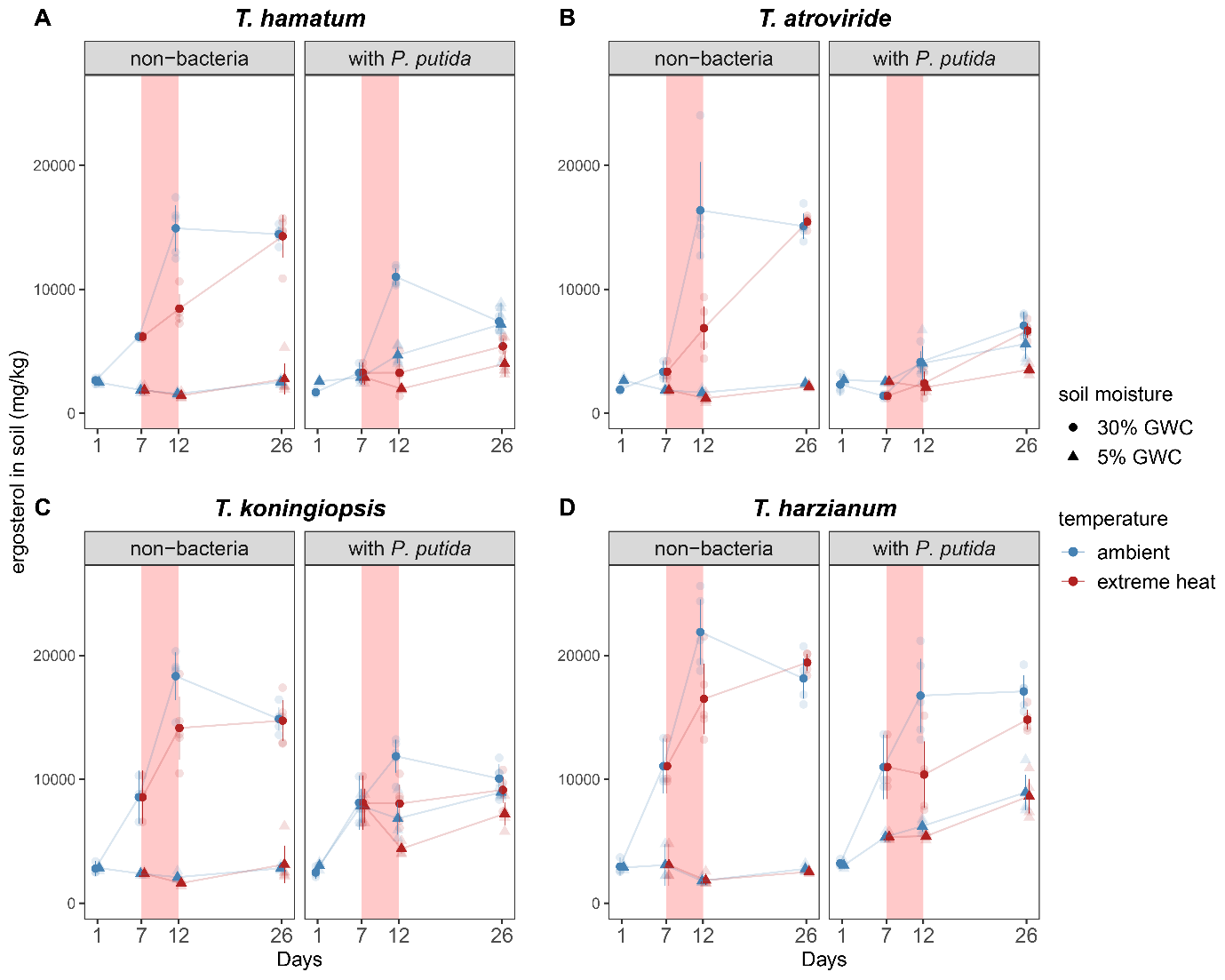
**Figure S3. Fungal biomass dynamics in soil.** The biomass (mean ± standard error) of *T. hamatum* (A), *T. atroviride* (B), *T. koningiopsis* (C), and *T. harzizanum* (D). n=3 for day 1 and 7, n = 5 for day 12 and 26. The red region between day 7 and day 12 indicates the period of extreme heat. GWC: Gravimetric water content.

**Table S1.** Comparison of candidate models for bacterial density (temperature, soil moisture, fungal presence, and time as the main effects).

| Linear model | df | AICc | ΔAICc | weight |
| --- | --- | --- | --- | --- |
| 2-way interactions with quadratic time | 16 | 26.673 | 0.000 | 1.000 |
| 2-way interactions with linear time | 12 | 96.000 | 69.328 | < 0.001 |
| Full interaction model with linear time | 17 | 98.922 | 72.250 | < 0.001 |

Note: df is the number of the estimated parameters in the model. AICc is the small-sample correction of AIC (Akaike Information Criterion), and a lower AICc means a better balance of fit and simplicity. ΔAICc represents how far each model is from the best model.

**Table S2.** Comparison of candidate models for bacterial density (temperature, soil moisture, fungal species, and time as the main effects).

| Linear model | df | AICc | ΔAICc | weight |
| --- | --- | --- | --- | --- |
| 2-way interactions with quadratic time | 31 | 18.968 | 0.000 | 1.000 |
| 2-way interactions with linear time | 24 | 94.122 | 75.154 | < 0.001 |
| Full interaction model with linear time | 41 | 118.170 | 99.202 | < 0.001 |

Note: df is the number of the estimated parameters in the model. AICc is the small-sample correction of AIC (Akaike Information Criterion), and a lower AICc means a better balance of fit and simplicity. ΔAICc represents how far each model is from the best model.

**Table S3.** Comparison of candidate models for fungal biomass across species (temperature, soil moisture, bacterial presence, and time as the main effects).

| Linear model | df | AICc | ΔAICc | weight |
| --- | --- | --- | --- | --- |
| 3-way interactions with quadratic time | 23 | 509.173 | 0.000 | 1.000 |
| Full interactions model with linear time | 17 | 641.601 | 132.428 | < 0.001 |
| 3-way interactions model with linear time | 16 | 642.208 | 133.036 | < 0.001 |

Note: df is the number of the estimated parameters in the model. AICc is the small-sample correction of AIC (Akaike Information Criterion), and a lower AICc means a better balance of fit and simplicity. ΔAICc represents how far each model is from the best model.

**Table S4.** Comparison of candidate models for fungal biomass (temperature, soil moisture, bacterial presence, fungal species, and time as the main effects).

| Linear model | df | AICc | ΔAICc | weight |
| --- | --- | --- | --- | --- |
| 3-way interactions with quadratic time | 68 | -9.253 | 0.000 | 1.000 |
| 3-way interactions model with linear time | 49 | 384.177 | 393.430 | < 0.001 |
| Full interactions model with linear time | 65 | 412.369 | 421.622 | < 0.001 |

Note: df is the number of the estimated parameters in the model. AICc is the small-sample correction of AIC (Akaike Information Criterion), and a lower AICc means a better balance of fit and simplicity. ΔAICc represents how far each model is from the best model.

**Table S5.** Type III ANOVA showing the effects of temperature, soil moisture, fungi, time, and their two-way interactions on bacterial density in soil.

| Factors | df | *F* value | *P* value |
| --- | --- | --- | --- |
| Temperature | 1 | **27.181** | ***P* < 0.001** |
| Soil moisture | 1 | **310.995** | ***P* < 0.001** |
| Fungi | 4 | **3.001** | **0.019** |
| Time (quadratic) | 2 | 0.188 | 0.829 |
| Temperature × Soil moisture | 1 | **25.861** | ***P* < 0.001** |
| Temperature × Fungi | 4 | **5.642** | ***P* < 0.001** |
| Temperature × Time (quadratic) | 2 | **18.705** | ***P* < 0.001** |
| Soil moisture × Fungi | 4 | 1.558 | 0.186 |
| Soil moisture × Time (quadratic) | 2 | **29.818** | ***P* < 0.001** |
| Fungi × Time (quadratic) | 8 | **2.574** | **0.011** |
| Residuals | 259 | NA | NA |

**Table S6.** Type III ANOVA showing the effects of temperature, soil moisture, bacteria, fungal species, time, and their three-way interactions on fungal biomass in soil.

| Factors | df | *F* value | *P* value |
| --- | --- | --- | --- |
| Temperature | 1 | **4.597** | **0.033** |
| Soil moisture | 1 | **524.009** | ***P* < 0.001** |
| Bacteria | 1 | **50.570** | ***P* < 0.001** |
| Fungal species | 3 | **16.049** | ***P* < 0.001** |
| Time (quadratic) | 2 | **109.196** | ***P* < 0.001** |
| Temperature × Soil moisture | 1 | **5.337** | **0.021** |
| Temperature × Bacteria | 1 | **13.003** | ***P* < 0.001** |
| Temperature × Fungal species | 3 | **2.849** | **0.037** |
| Temperature × Time (quadratic) | 2 | **9.844** | ***P* < 0.001** |
| Soil moisture × Bacteria | 1 | **219.064** | ***P* < 0.001** |
| Soil moisture × Fungal species | 3 | **4.641** | **0.003** |
| Soil moisture × Time (quadratic) | 2 | **103.095** | ***P* < 0.001** |
| Bacteria × Fungal species | 3 | **18.347** | ***P* < 0.001** |
| Bacteria × Time (quadratic) | 2 | 1.477 | 0.230 |
| Fungal species × Time (quadratic) | 6 | **5.813** | ***P* < 0.001** |
| Temperature × Soil moisture × Bacteria | 1 | 1.465 | 0.227 |
| Temperature × Soil moisture × Fungal species | 3 | 0.557 | 0.644 |
| Temperature × Soil moisture × Time (quadratic) | 2 | **6.260** | **0.002** |
| Temperature × Bacteria × Fungal species | 3 | **4.680** | **0.003** |
| Temperature × Bacteria × Time (quadratic) | 2 | **4.229** | **0.015** |
| Temperature × Fungal species × Time (quadratic) | 6 | 2.115 | 0.051 |
| Soil moisture × Bacteria × Fungal species | 3 | 2.113 | 0.098 |
| Soil moisture × Bacteria × Time (quadratic) | 2 | **59.349** | ***P* < 0.001** |
| Soil moisture × Fungal species × Time (quadratic) | 6 | **7.075** | ***P* < 0.001** |
| Bacteria × Fungal species × Time (quadratic) | 6 | **3.439** | **0.003** |
| Residuals | 397 | NA | NA |

**Sample preparation for microscopic imaging**

For each fungal species, an agar plug with fungal hyphae (diameter: 5 mm) was placed in the centre of the PAD plate (diameter: 9 cm), and the bacteria were added in droplets (OD600 = 0.1, 1 µL/droplet) at 1 cm from the fungal plug. We then incubated the plates at 28°C upright. After 24 h of incubation, we observed and captured images of the bacterial colonies on the fungal hyphae using a Nikon SMZ25 stereomicroscope with SHR Plan Apo 0.5× objective, in both the brightfield and GFP channels (excitation: 460-500 nm, emission: 510-560 nm). We then cut out the bacterial colonies with fungal hyphae from the plate with a scalpel (sample size: approximately 7 mm × 7 mm) and placed the samples in fixative solution (2.5% Glutaraldehyde in 0.15 M Hepes). The samples were then dried in a critical-point drying apparatus in CO_2_ and then coated with gold for scanning electron microscopy (Zeiss EVO 40).
